# Longitudinal Dynamics and Site-Specific Recovery of the Human Respiratory Microbiome Following Smoking Cessation

**DOI:** 10.64898/2026.02.03.702818

**Authors:** Silvia Gschwendtner, Draginja Kovacevic, Karoline I. Gaede, Christian Herzmann, Jörg Overmann, Michael Schloter, Susanne Krauss-Etschmann

## Abstract

**Background:** The human respiratory tract harbours diverse microbial communities crucial for health, but their dynamics during environmental perturbations like smoking remain poorly understood. While smoking is a major risk factor for various diseases, its compartment-specific effects on the respiratory microbiome and potential recovery following cessation have not been fully elucidated. Here, we present a longitudinal, multi-site study of respiratory microbiome dynamics in smokers undergoing cessation, benchmarked against healthy never-smokers.

**Methods:** Using standardized sampling of the anterior nares, oropharynx, and bronchoalveolar lavage (BAL), combined with 16S rRNA gene amplicon sequencing and rigorous contamination controls, we characterized community composition, diversity, personalization, and microbial interactions across airway compartments.

**Results:** Smokers exhibited pronounced microbiome alterations: nasal richness increased, while lung richness and core taxa were reduced. Smoking-induced changes were compartment-specific and most pronounced in nose and lung. The degree of individuum-specific differences in community structure was elevated in smokers and correlated with smoking intensity and duration. Short-term cessation (6 weeks) led to minor shifts in taxa abundance but increased similarity between oropharyngeal and lung communities, whereas long-term cessation (1 year) resulted in partial restoration, particularly in lung and nasal microbiomes. Some taxa, including *Haemophilus* and *Prevotella_7*, showed persistent alterations, highlighting lasting smoking effects. Network analyses revealed that smoking disrupted microbial co-occurrence and reduced community connectivity, whereas cessation partially restored interaction networks, with dynamics differing between oropharynx and lung, reflecting different underlying ecological assembly processes. Recovery trajectories were highly individualized, with lung microbiomes influenced by deterministic processes and upper airway microbiomes shaped by stochastic factors, explaining site-specific responses and the persistence of personalized microbial signatures.

**Conclusion:** These results provide the first time-resolved, multi-compartment characterization of respiratory microbiome recovery after smoking cessation, revealing that smoking leaves long-lasting, site-specific imprints on airway microbial communities and interactions. Our findings underscore the need for individual and compartment-specific approaches when designing microbiome-based interventions to support respiratory health.

## Background

Human-associated microbial communities have co-evolved with their host over millions of years, colonizing nearly all body surfaces, including the respiratory tract (RT). The RT comprises the upper respiratory tract (URT; nose, sinuses, naso- and oropharynx, supraglottic larynx) and the lower respiratory tract (LRT; infraglottic larynx, trachea, lungs). In adults, the airways cover ∼70 m² (≈40 times the skin surface; (^1^)) and are populated by compartment-specific microbiota essential for RT function and host health. Community composition of the RT reflects a dynamic balance between immigration (from the oral cavity and URT), elimination (via mucociliary clearance, cough, and immunity), and differential growth (^2, 3^).

The anterior nares, lined by keratinized epithelium, harbor typical skin colonizers (*Corynebacterium*, *Propionibacterium*, *Staphylococcus*) alongside *Dolosigranulum*, *Moraxella*, and *Streptococcus* (^4^). The nasopharynx contains overlapping but more diverse taxa, including *Haemophilus*, whereas the oropharynx is enriched in *Streptococcus*, *Neisseria*, *Rothia*, and anaerobes such as *Leptotrichia*, *Prevotella* and *Veillonella*. A balanced URT microbiome is critical for pathogen exclusion: *Dolosigranulum* and *Corynebacterium* are linked to reduced *Streptococcus pneumoniae* colonization (^5, 6^), and *Staphylococcus epidermidis* can inhibit *S. aureus* biofilms (^7^).

The LRT was long considered sterile; however molecular analyses have revealed that healthy lungs harbor low biomass yet diverse microbial communities in healthy lungs, dominated by *Prevotella*, *Streptococcus*, *Veillonella*, *Actinomyces*, *Fusobacterium*, and *Haemophilus* (^8–12^). This discovery has fundamentally reshaped our understanding of respiratory biology, highlighting the microbiota as a key determinant of pulmonary health and disease.

Smoking is a major risk factor for lung cancer, chronic obstructive pulmonary disease (COPD), cardiovascular disease, and periodontitis (^13^). Increasing evidence suggests that disease initiation and progression may be partly mediated by smoking-induced dysbiosis of the respiratory microbiome (^14^). Proposed mechanisms include altered oxygen tension, immune suppression, promotion of biofilm formation, and direct exposure to smoke-derived chemicals or bacteria present in cigarettes (^14–17^).

Despite numerous investigations, the impact of smoking on respiratory microbiota remains inconsistent and highly site dependent. Some studies report increased microbial diversity in the oral cavity and URT (^11, 18–20^), while others observe reduced diversity (^21, 22^). In contrast, reduced diversity is more frequently observed in the lower airways (^11, 23^). Although smoking-induced shifts in community composition are commonly detected, the specific taxa involved vary considerably, limiting their predictive value.

Notably, some of these smoking-induced alterations appear reversible. In mice, oropharyngeal microbial diversity and composition normalized within three months of smoking cessation (^24^). Comparable recovery was observed in the human nasopharynx after 12–15 months (^25^). However, persistent changes in the LRT (^23^) suggest that smoking exerts lasting, compartment-specific effects on the airway microbiome, even in otherwise healthy individuals.

Despite accumulating evidence that a (partial) reestablishment of a health microbiome of smokers might be possible after smoking cessation, the temporal dynamics of respiratory microbiome recovery at different compartments of the RT as well as personalized pattern are still not well understood.

Here, we present longitudinal microbiome profiles from multiple respiratory sites during smoking cessation, benchmarked against healthy never-smokers. This dataset enables, for the first time, a time-resolved characterization of microbiome regeneration across the respiratory tract and reveals inter-individual variation in recovery trajectories. Our findings provide a foundation for targeted microbiome-based interventions, particularly during the early phases of smoking cessation when communities are most perturbed and only partially restored.

## Materials and Methods

### Participant selection and sampling

The LuMEn cohort was recruited at the Center for Clinical Studies, Research Center Borstel, Germany, between March 2017 and July 2020 as described previously (^12^). Eligible participants were Caucasian adults (≥18 years) who met the following criteria: absence of (1) respiratory infection or systemic glucocorticoid therapy within the past month, (2) antibiotic therapy within the past two months, (3) diabetes mellitus, (4) pregnancy or lactation, (5) active or previous tuberculosis, (6) immunosuppression, or (7) known pulmonary disease, except for COPD stage GOLD I/II. Individuals with medical contraindications to bronchoscopy, as determined by the investigator (e.g., allergy to sedative agents), were excluded. Demographic characteristics are presented in Table 1. Medical history, environmental exposures, and socioeconomic status were recorded using a standardized questionnaire. Based on CDC definitions, participants were categorized as active smokers (AS), former smokers (FS), or never-smokers (NS).

**Table 1.** Summary of baseline characteristics of the study cohort.

|  | average | sd | se |
| --- | --- | --- | --- |
| **number, n** | 25 |  |  |
| age in years | 38.96 | 12.8 | 2.56 |
| male, n (%) | 14 | (56.0%) |  |
| height | 175.68 | 11.28 | 2.26 |
| weight | 81.48 | 16.35 | 3.27 |
| **never-smokers, n (%)** | 10 | (40.0) |  |
| age in years | 37.4 | 13.58 | 4.3 |
| male, n (%) | 4 | (40.0%) |  |
| height | 172.4 | 10.38 | 3.28 |
| weight | 82.1 | 10.83 | 3.42 |
| nicotine | 2.3 | 0.67 | 0.21 |
| cotinine | 5.3 | 9.75 | 3.08 |
| anabasine | 2 | 0 | 0 |
| OH_cotinine | 108.1 | 215.35 | 68.1 |
| **active smokers, n (%)** | 15 | (60.0) |  |
| age in years | 40 | 12.62 | 3.26 |
| male, n (%) | 10 | (66.7%) |  |
| height | 177.87 | 11.65 | 3.01 |
| weight | 81.07 | 19.55 | 5.05 |
| nicotine | 447.13 | 550.1 | 142.04 |
| cotinine | 1052.73 | 632.81 | 163.39 |
| anabasine | 6.73 | 4.7 | 1.21 |
| OH_cotinine | 5075.4 | 5357.21 | 1383.23 |
| smoking years | 24 | 12.74 | 3.29 |
| cpd | 14.73 | 5.12 | 1.32 |
| cpd_max | 29.2 | 8.8 | 2.27 |
| pack years | 21.13 | 18.93 | 4.89 |
| **ex-smokers 6w, n (%)** | 15 | (60.0) |  |
| age in years | 40 | 12.62 | 3.26 |
| male, n (%) | 10 | (66.7%) |  |
| height | 177.87 | 11.65 | 3.01 |
| weight | 81.07 | 19.55 | 5.05 |
| nicotine | 3 | 2.1 | 0.54 |
| cotinine | 7.87 | 14.16 | 3.66 |
| anabasine | 2 | 0 | 0 |
| OH_cotinine | 64.73 | 51.87 | 13.39 |
| smoking years | 24 | 12.74 | 3.29 |
| cpd | 14.73 | 5.12 | 1.32 |
| cpd_max | 29.2 | 8.8 | 2.27 |
| pack years | 21.13 | 18.93 | 4.89 |
| **ex-smokers 1yr, n (%)** | 5 | (20.0) |  |
| age in years | 36.6 | 14.69 | 6.57 |
| male, n (%) | 3 | (60.0%) |  |
| height | 172.4 | 11.84 | 5.3 |
| weight | 84.14 | 27.75 | 12.41 |
| nicotine | 2 | 0 | 0 |
| cotinine | 2 | 0 | 0 |
| anabasine | 2 | 0 | 0 |
| OH_cotinine | 40 | 0 | 0 |
| smoking years | 20.6 | 14.79 | 6.62 |
| cpd | 15.8 | 6.42 | 2.87 |
| cpd_max | 28.6 | 9.48 | 4.24 |
| pack years | 21.2 | 27.07 | 12.11 |

The present study sub-cohort involved of 25 subjects (11 females and 14 males), including 10 NS and 15 AS. Samples after cessation were taken at 6 weeks (FS6w; 15 subjects) and 1 year (FS1yr; 5 subjects because only those completed 1 year cessation). Age distribution was comparable between NS and AS (p=0.689), while NS had higher number of females compared to AS (p=0.007) (n=25; Table 1). Nicotine, cotinine, anabasine, and OH-cotinine concentrations were quantified in unprocessed urine samples using liquid chromatography-tandem mass spectrometry (LC-MS/MS). Quality controlled biosampling included deep bilateral nasal swabs (n=36), bilateral oropharyngeal swabs (n=45), and bronchoalveolar lavage (BAL; n=45) representing both important URT and LRT compartments. Nasal and oropharyngeal swabs (Mast swab TM, Mast Group Ltd., UK) were immediately frozen in cryotubes on dry ice. Flexible bronchoscopy was performed in accordance with German guidelines. The bronchoscope was advanced into a subsegmental bronchus of the middle lobe, and BAL was carried out with 15 aliquots of 20 mL sterile saline (0.9%), for a total of 300 mL. The first BAL fraction was discarded due to possible contamination, while the remaining fractions were pooled and stored at –80 °C. Sterile saline solution was included as a control for microbiome analyses.

The study was approved by the Ethics Committee of the University of Luebeck (ref. 16-145) and registered at clinicaltrials.gov (NCT03562442; first submitted: May 26, 2018). Oral and written informed consent was obtained from all participants in accordance with ICH/GCP guidelines.

### DNA extraction

The cell pellet of 5 mL BAL (centrifuged at 20.000 x g/10 min) was dissolved in 180 µl lysozyme solution (10mg/µl), whereas nasal and oropharyngeal swabs were covered directly with 400 µl lysozyme solution (10mg/µl). After incubation for 45 min at 37°C, DNA extraction was performed using the PureLink™ Genomic DNA Mini Kit (ThermoFisher Scientific, Altham, USA) according to the manufacturer’s protocol for Gram-positive bacteria. To identify contaminants deriving from sampling and extraction kit, controls were included (DNA extraction of 5 mL sterile saline solution (8 samples) and 5 mL bronchoscope flushing (8 samples), as well as 5 blank extractions).

### 16S rRNA gene library preparation

Amplicon sequencing of the V4 hypervariable region of the 16S rRNA gene was performed on a MiSeq Illumina instrument (MiSeq Reagent Kit v3 (600 Cycle); Illumina, San Diego, CA, USA) using the universal eubacterial primers 515F (^26^) and 806R (^27^). PCR was done using NEBNext high fidelity polymerase (New England Biolabs, Ipswich, USA) in a total volume of 25 µl (10 ng DNA template, 12.5 µl polymerase, 5 pmol of each primer, 2.5 µl 3% BSA) and the following PCR conditions: 5 min at 98°C; 30 cycles of 10 s at 98°C, 30 s at 55 °C, 30 s at 72 °C; 5 min 72 °C. To identify potential contaminants deriving from library preparation, we included 4 blank PCR controls. Subsequent library preparation was performed as described previously (^28^). The sequence data generated in this study have been deposited in the NCBI Short Read Archive (SRA) under accession number PRJNA1328433 and will be made publicly available upon acceptance of the manuscript.

### Sequence processing

FASTQ files were trimmed with a minimum read length of 50 using Cutadapt (^29^) and quality control was performed via FastQC (^30^). For subsequent data analysis, the DADA2 pipeline v 1.30 (^31^) was used with the following trimming and filtering parameters: 20 bp were removed n-terminally and reads were truncated at position 260 (forward) and 200 (reverse), respectively, with expected error of 4 (forward) and 6 (reverse). Taxonomic analysis was performed using SILVA v138.1. Reads were excluded if classified as mitochondria or chloroplast or if the phylum was missing. All blank samples (saline solution, bronchoscope flushing, blank extraction and PCR no template control) were analyzed together with biological samples and showed clearly lower richness and different beta diversity compared to respiratory tract samples (Additional Figure 1). Sequences present in at least 50% of blanks were considered as core contaminome. Additionally, potential contaminants were identified statistically via the ‘decontam’ R package (^32^). All sequences of the core contaminome and/or being identified by ‘decontam’ were removed from sample data (in total 45, Additional Table 1), resulting in a total amount of 5,446,191 reads (corresponding to an average of 43,223 reads per sample) assigned to 2,221 amplicon sequence variants (ASV).

**Figure 1.**
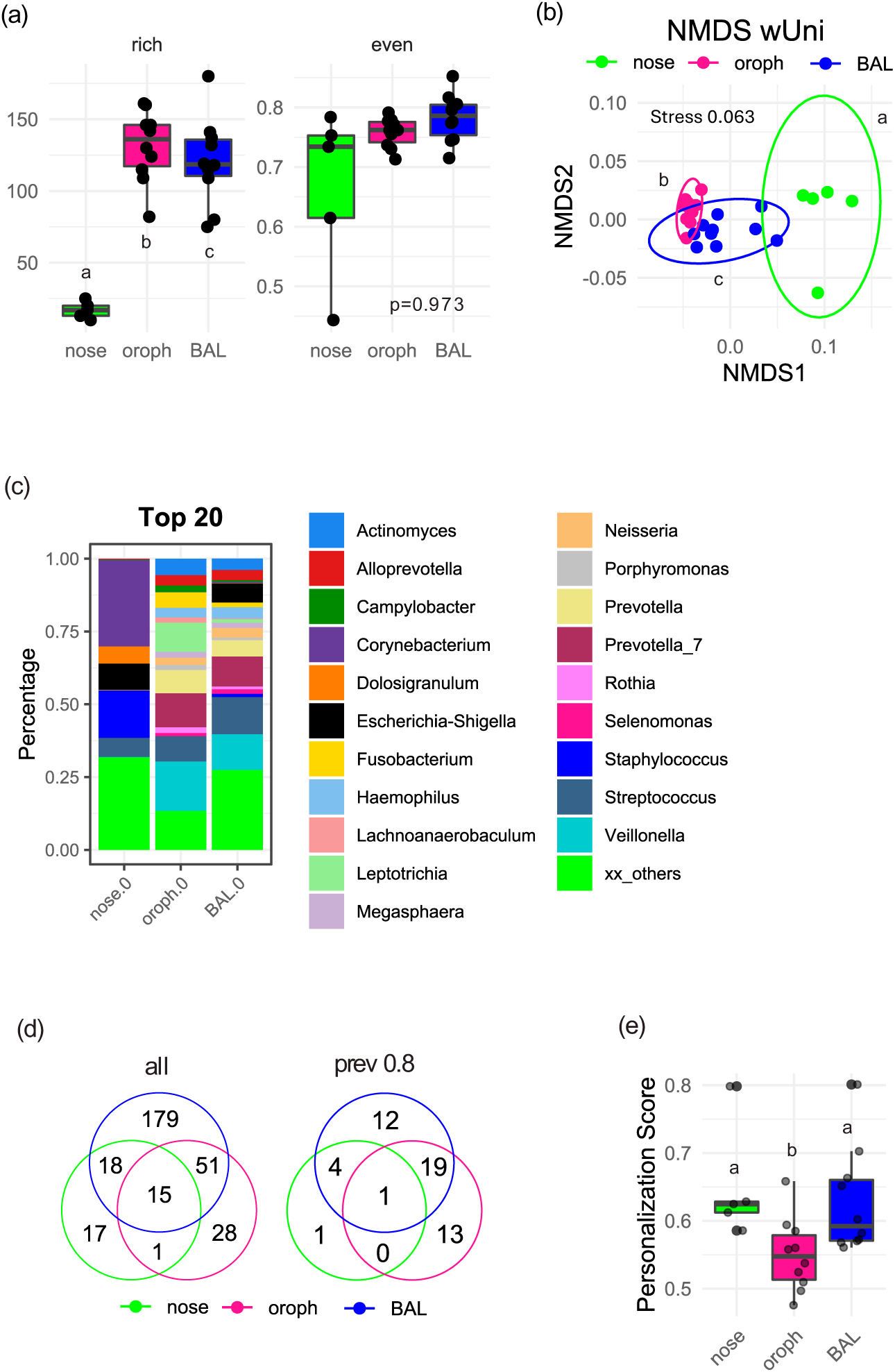
Microbial communities among the respiratory tract of never-smokers (NS) (a) Boxplots of alpha diversity measures showed decreasing richness oropharynx>BAL>nose, while evenness did not differ. (b) NMDS plot of weighted UniFrac distances revealed distinct microbial communities in all airway cavities. Letters in (a, b) indicate significant differences (p<0.05) calculated via generalized linear mixed-effect models including the study subjects as random effect term and PERMANOVA using strata to restrict permutations within blocks with Benjamini-Hochberg correction for multiple comparisons. (c) Relative abundance of the top 20 genera showed overlap in abundant genera between oropharynx and BAL, while nose clearly differed. (d) Venn diagram showed number of shared and unique genera for nose, oropharynx and BAL samples (left: without prevalence threshold, right: prevalence 0.8). (e) Personalization scores revealed comparable individual variability in nose and BAL and lower in oropharynx. Letters indicate significant differences (p<0.05) calculated via linear mixed-effect models including the study subjects as random effect with Benjamini-Hochberg correction for multiple comparisons.

### Statistical analysis

All plots and statistics were performed in R version 4.4.0 (https://www.R-project.org). Sequencing data were normalized using cumulative-sum scaling (CSS) via the metagenomeSeq R package (^33^). CSS calculates scaling factors based on the cumulative sum of gene abundances up to a dataset-specific threshold, allowing an accurate comparison of microbial communities across different samples with varying sizes. Alpha diversity was calculated using species richness, evenness as well as Shannon and Simpson diversity index. Beta diversity was analysed via unweighted and weighted UniFrac distance as well as Bray-Curtis dissimilarity. For statistical purposes, generalized linear mixed-effects models including the study subjects (longitudinal samples) as repeated measurements by defining a random effect term (R package lme4) and PERMANOVA using strata to restrict permutations within blocks (R package vegan) were used. P value adjustment for multiple comparisons was performed with Benjamini-Hochberg correction.

To identify microbial taxa differing between never-smokers (NS), active smokers (AS) and former smokers (FS), DESeq2 was used (^34^). Taxa were considered as enriched or depleted with a log2-fold change >=2 and an FDR-adjusted p-value < 0.05.

For analysis of individual variability, a personalization score for each subject was calculated as difference between inter- and intra-individual distances based on Bray-Curtis dissimilarity measures. In addition, Bray-Curtis distances were used to assess the recovery potential after cessation by computing the similarity of the microbial community of former smokers per subject towards the NS community. Furthermore, the similarity of oropharynx and BAL microbial communities within each subject was computed via between-group distances calculated by weighted UniFrac distance measures. Statistical analysis was performed using linear mixed-effects models including the study subjects (longitudinal samples) as repeated measurements by defining a random effect term (R package lme4) and P value adjustment for multiple comparisons with Benjamini-Hochberg correction. To search for correlations between smoking-related factors (smoking years, cigarettes per day (cpd), packyears and exposure based on nicotine levels) and personalization scores or recovery, Pearson correlations were calculated, using Benjamini-Hochberg p value adjustment for multiple pairwise comparisons. To find correlations between numeric variables and microbiome data, Spearman correlations were calculated between clr-transformed abundance data and meta variables, using Benjamini-Hochberg p value adjustment for multiple pairwise comparisons. The core microbiome per group (respiratory cavity, smoking status) was defined as genera present in at least 80 % of subjects without setting an additional abundance cutoff.

For the determination of microbial assembly processes, the ßNTI and Raup-Crick-based Bray-Curtis (RCbray) method described by Stegen et al. (^35^) was used. Briefly, deterministic selection (|bNTI|>2) can be distinguished into heterogeneous selection (selective pressures drive communities to divergent configurations) and homogeneous selection (selective pressures push communities towards a common composition), whereas stochastic processes (|bNTI|<2) are classified into homogenizing dispersal (communities are more similar than expected; populations are capable of interactions, allowing members to freely exchange), dispersal limitation (populations are unable to mix leading to development via ecological drift) and undominated processes (no assembly process is capable of explaining variation) according to RCbray values.

Microbial co-occurrence networks were inferred via SparCC correlations implemented in the R package NetCoMi v1.1.0 (36), using the “signed” transformation to transform the estimated associations into dissimilarities and including only ASV present in 50% of the samples per group. Highly interconnected modules of nodes within the networks were then identified using the cluster_fast_greedy algorithm. All plots were created in R using ggplot2 (^37^), ggpubr (^38^) and microViz (^39^).

## Results

### Bacterial community composition among the respiratory tract in never-smokers (NS)

Microbial diversity varied across respiratory sites, with richness decreasing from oropharynx to lung to nose, while evenness remained similar (Fig. 1a). Beta diversity confirmed distinct, compartment-specific communities (Fig. 1b). Nasal samples were dominated by *Corynebacterium, Dolosigranulum, Escherichia-Shigella, Streptococcus* and *Staphylococcus*, whereas oropharynx and BAL shared a core microbiome dominated by *Actinomyces*, *Prevotella* groups, *Streptococcus*, and *Veillonella*, with *Fusobacterium* and *Leptotrichia* enriched in the oropharynx and *Escherichia-Shigella* in BAL (Fig. 1c). Richness on the level of bacterial genera was highest in BAL (263), followed by the oropharynx (95) and the nose (51), with many taxa being unique to each site (Fig. 1d; Additional Table 2). Core microbiome analysis (≥ 80% prevalence) revealed limited overlap across compartments: only *Streptococcus* was shared by all, while oropharynx and BAL showed the strongest convergence. The marked reduction from total to core genera indicated substantial inter-individual variability.

**Table 2.** Topological features of bacterial co-occurrence network analysis including number of nodes, clustering coefficient, modularity, positive edge percentage (PEP), edge density, robustness, average path length, centrality measures (degree, between, close, eigenvector) and number of modules.

| oroph |  |  |  |  |
| --- | --- | --- | --- | --- |
|  | NS | AS | FS_6w | FS_1yr |
| **Nodes** | 89 | 85 | 87 | 110 |
| **Clustering coef.** | 0.54 | 0.48 | 0.48 | 0.70 |
| **Modularity** | 0.05 | 0.09 | 0.08 | 0.02 |
| **PEP** | 50.74 | 50.59 | 51.07 | 49.01 |
| **Edge density** | 0.45 | 0.35 | 0.36 | 0.64 |
| **Robustness** | 0.13 | 0.09 | 0.09 | 0.21 |
| **Av. path length** | 0.98 | 1.06 | 1.05 | 0.90 |
| **degree** | 0.45 | 0.35 | 0.36 | 0.65 |
| **between** | 0.01 | 0.01 | 0.01 | 0.00 |
| **close** | 1.13 | 1.04 | 1.05 | 1.37 |
| **eigen** | 0.59 | 0.55 | 0.60 | 0.64 |
| **Module no.** | 3 | 4 | 4 | 3 |
| **Hubs** | Asv1;Veillonella | Asv109;Prevotella_7 | Asv1;Veillonella | Asv110;Mogibacterium |
|  | Asv13;Streptococcus | Asv16;Prevotella | Asv35;Veillonella | Asv16;Prevotella |
|  | Asv4;Streptococcus | Asv35;Veillonella | Asv45;Stomatobaculum | Asv26;Selenomonas |
|  | Asv43;Streptococcus | Asv45;Stomatobaculum | Asv82;Oribacterium | Asv356;Treponema |
|  | Asv44;Gemella | Asv9;Megasphaera | Asv9;Megasphaera | Asv56;Leptotrichia |
|  |  |  |  | Asv9;Megasphaera |
| BAL |  |  |  |  |
|  | NS | AS | FS_6w | FS_1yr |
| **Nodes** | 66 | 21 | 43 | 70 |
| **Clustering coef.** | 0.77 | 0.35 | 0.41 | 0.68 |
| **Modularity** | -0.01 | 0.21 | 0.13 | 0.02 |
| **PEP** | 49.03 | 52.33 | 51.27 | 48.99 |
| **Edge density** | 0.65 | 0.21 | 0.29 | 0.64 |
| **Robustness** | 0.20 | 0.06 | 0.06 | 0.18 |
| **Av. path length** | 1.01 | 1.43 | 1.16 | 0.93 |
| **degree** | 0.66 | 0.24 | 0.29 | 0.64 |
| **between** | 0.01 | 0.04 | 0.02 | 0.01 |
| **close** | 1.15 | 0.85 | 0.96 | 1.26 |
| **eigen** | 0.67 | 0.49 | 0.56 | 0.64 |
| **Module no.** | 2 | 5 | 6 | 3 |
| **Hubs** | Asv15;Alloprevotella | Asv4;Streptococcus | Asv12;Prevotella_7 | Asv13;Streptococcus |
|  | Asv2;Prevotella_7 | Asv44;Gemella | Asv14;Neisseria | Asv18;Actinomyces |
|  | Asv21;Campylobacter |  | Asv47;Staphylococcus | Asv26;Selenomonas |
|  | Asv7;Fusobacterium |  |  | Asv3;Prevotella_7 |

Personalization scores (Bray–Curtis) were high in the nose and BAL, but significantly lower in the oropharynx (Fig. 1e), with no correlation to age (p=0.537) or sex (p=0.726) (data not shown).

### Bacterial community composition among the respiratory tract in active smokers (AS)

As for NS, all compartments of AS differed in microbial diversity and community composition (Fig. 2a,b), with richness being reduced in BAL (AS: 81 vs. NS: 120, p=0.041), increased in the nose (AS: 29 vs. NS: 17, p=0.009), and unchanged in the oropharynx (Fig. 3a). The nasal microbiome of AS was primarily composed of *Corynebacterium*, *Dolosigranulum, Prevotella, Staphylococcus*, *Streptococcus* and *Veillonella* (Fig. 2c), while the oropharynx and BAL were both dominated by *Actinomyces*, *Prevotella*, *Streptococcus*, and *Veillonella*, with *Fusobacterium* and *Leptotrichia* also being abundant in the oropharynx.

**Figure 2.**
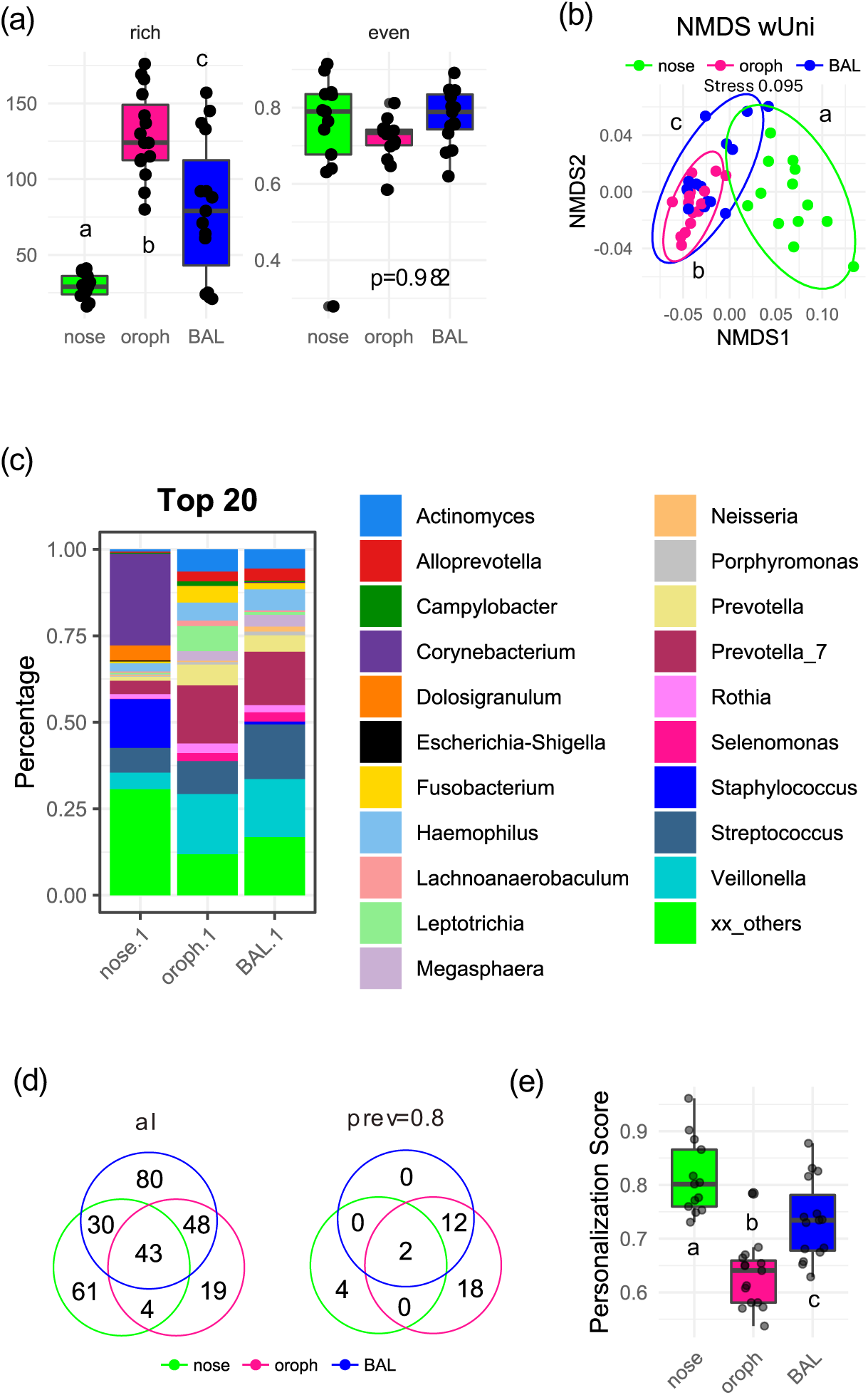
Microbial communities among the respiratory tract of active smokers (AS) (a) Boxplots of alpha diversity measures showed decreasing richness oropharynx>BAL>nose, while evenness did not differ. (b) NMDS plot of weighted UniFrac distances revealed distinct microbial communities in all airway cavities. Letters in (a, b) indicate significant differences (p<0.05) calculated via generalized linear mixed-effect models including the study subjects as random effect term and PERMANOVA using strata to restrict permutations within blocks with Benjamini-Hochberg correction for multiple comparisons. (c) Relative abundance of the top 20 genera showed overlap in abundant genera between oropharynx and BAL, while nose clearly differed. (d) Venn diagram showed number of shared and unique genera for nose, oropharynx and BAL samples (left: without prevalence threshold, right: prevalence 0.8). (e) Personalization scores revealed decreasing individual variability nose>BAL> oropharynx. Letters indicate significant differences (p<0.05) calculated via linear mixed-effect models including the study subjects as random effect with Benjamini-Hochberg correction for multiple comparisons.

**Figure 3.**
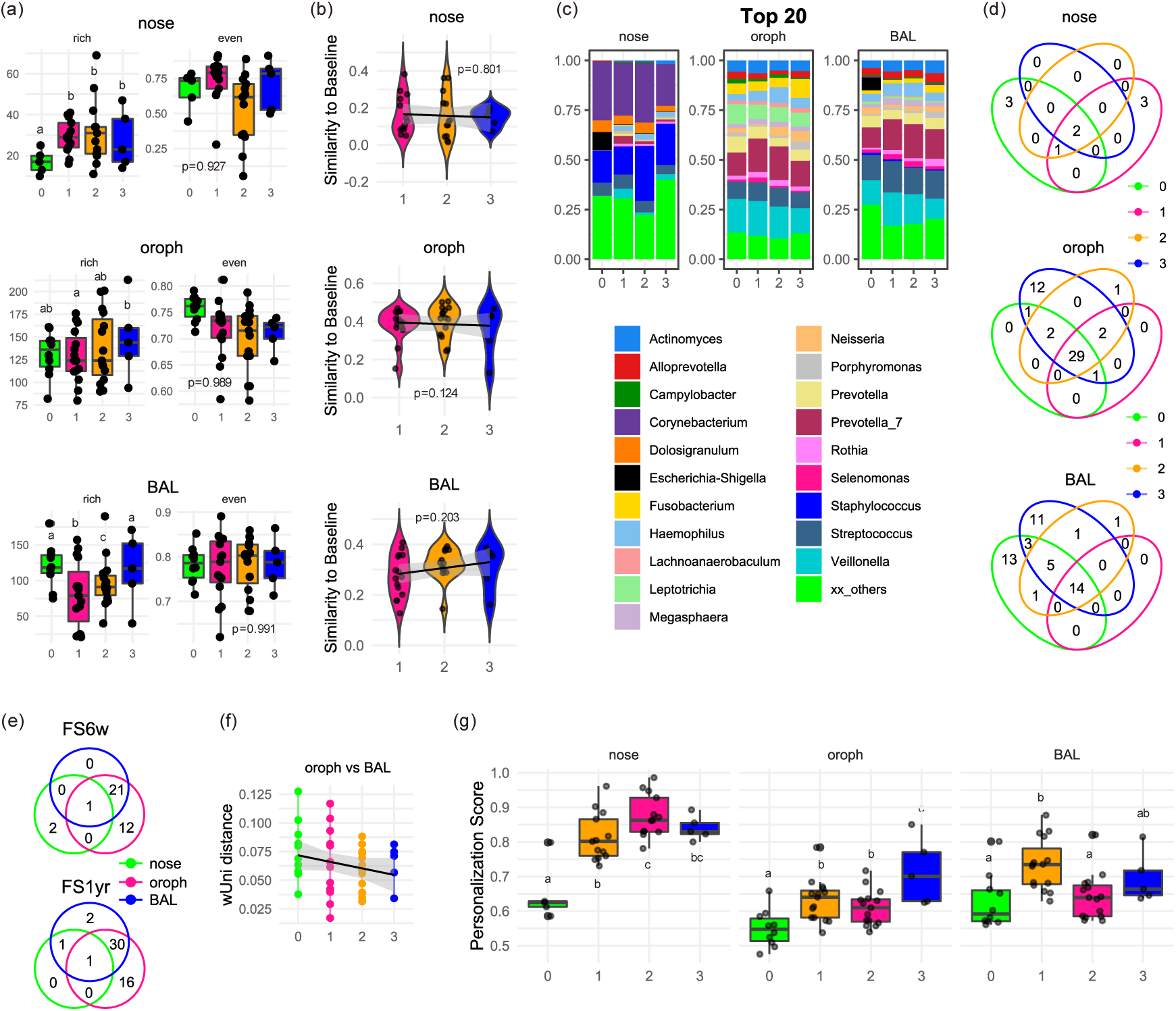
Response to smoking and cessation (a) Boxplots of alpha diversity measures among the different smokestatus (0: NS, 1: AS, 2: FS6w, 3: FS1yr) revealed a highly cavity-dependent response to smoking and cessation, with most pronounced effects on nose (lasting increase of diversity during smoking) and BAL (reduction of diversity during smoking but recovery after cessation). Whereas smoking did not affect the diversity in oropharynx, richness increased after long-term cessation. Letters indicate significant differences (p<0.05) calculated via generalized linear mixed-effect models including the study subjects as random effect term, and Benjamini-Hochberg correction for P value adjustment. (b) Recovery trajectory measured as similarity of the microbial community of former smokers towards the NS community (baseline) for the different smokestatus (1: AS, 2: FS6w, 3: FS1yr) revealed no significant recovery of community composition after cessation. (c) Relative abundance of the top 20 genera showed site-specific changes for the different smokestatus ((0: NS, 1: AS, 2: FS6w, 3: FS1yr) in nose, oropharynx and BAL. (d) Venn diagrams showed the number of shared and unique core taxa (defined as genera present in at least 80% of subjects per group) between the different smokestatus (0: NS, 1: AS, 2: FS6w, 3: FS1yr) for nose, oropharynx and BAL. (e) Venn diagrams showed number of shared and unique core taxa (defined as genera present in at least 80% of subjects per group) for nose, oropharynx and BAL samples after short-(FS6w) and long-term cessation (FS1yr). (f) Comparison of oropharyngeal and BAL communities within subjects at different smokestatus (0: NS, 1: AS, 2: FS6w, 3: FS1yr) showed a higher similarity between oropharynx and BAL after cessation compared to NS. (g) Personalization scores revealed a long-lasting increase of individual variability during smoking, with site-dependent intensity and most pronounced in nose. Letters indicate significant differences (p<0.05) calculated via linear mixed-effect models including the study subjects as random effect term, and Benjamini-Hochberg correction for P value adjustment.

Relative to NS, AS showed higher abundances of *Haemophilus*, *Mycoplasma*, *Prevotella* groups and *Veillonella* but reduced *Escherichia-Shigella* in the nose, altered levels of *Candidatus Saccharimonas*, *Mogibacterium*, *Neisseria*, *Prevotella_7* and *Selemonas* in the oropharynx, and a loss of *Escherichia-Shigella* with shifts in low-abundance taxa (e.g., reduced *Alistipes*, *Bacteroides*, *Muribaculaceae*, *Prevotellaceae* UCG-001; increased *Filifactor*, *Treponema*) in BAL (Fig. 3c, Additional Figure 2).

In AS, 138, 114, and 201 genera were observed in the nose, the oropharynx, and BAL, respectively, with 43 being common across the sites (Fig. 2d, Additional Table 2). Core microbiota (≥ 80% prevalence) were reduced compared to NS (nose: 6, oropharynx: 32, BAL: 14), with only *Streptococcus* and *Veillonella* being shared across all. No BAL-specific core genera were detected, in contrast to NS. Smoking thus reshaped the microbiota throughout the RT, most pronounced in the nose (increased richness and unique genera) and BAL (decreased richness and loss of core taxa). Bray–Curtis distances highlighted decreasing similarity between NS and AS communities from oropharynx (0.39) to BAL (0.28) to nose (0.17) (Fig. 3b). Smoking also increased personalization scores at all sites (Fig. 3g), with inter-individual variability highest in nose, intermediate in BAL, and lowest in oropharynx (Fig. 2e). Notably, variability was positively correlated with smoking years and pack-years in oropharynx and BAL, but not in nose (Additional Figure 3).

### Response to short-term smoking cessation is mediated by respiratory tract cavity

After 6 weeks of smoking cessation (FS6w), the richness of the nasal microbiome remained elevated compared to NS, whereas BAL richness partially recovered, and oropharyngeal richness was unchanged (Fig. 3a). Recovery trajectories based on Bray–Curtis distances showed no significant community-level recovery (Fig. 3b). Recovery potential in oropharynx and BAL correlated negatively with smoking duration, pack-years, and cigarette load, while in nose only maximum cigarettes/day was associated (Additional Figure 3).

At the genus level, only modest shifts were observed: in the nose, *Prevotella* and *Veillonella* decreased towards NS levels, whereas *Haemophilus*, *Mycoplasma*, and *Prevotella_7* remained elevated (Fig. 3c, Additional Figure 2). In the oropharynx, *Candidatus Saccharimonas*, *Neisseria* and *Selemonas* normalized, while *Mogibacterium* and *Prevotella_7* remained altered and *Haemophilus* increased. In BAL, only *Filifactor* decreased, whereas several low-abundance taxa (*Alistipes*, *Bacteroides*, *Muribaculaceae*, *Prevotellaceae* UCG-001) remained depleted.

Core microbiota analysis identified 3, 34, and 22 genera for the nose, oropharynx, and BAL, respectively (Fig. 3e, Additional Table 2). Shared genera between FS6w and NS but absent in AS (e.g., *[Eubacterium] nodatum* group, *Bergeyella* in oropharynx; *Campylobacter*, *Neisseria*, *Selenomonas* in BAL) may represent early markers of short-term microbial recovery following smoking cessation (Fig. 3d, Additional Table 2). Like in AS, BAL lacked unique core genera but overlap with oropharynx increased (21 vs. 12 genera in AS), suggesting convergence after cessation (Fig. 3e). Weighted UniFrac distances confirmed greater oropharynx–BAL similarity in FS6w vs. NS (p=0.046, Fig. 3f). Positive correlations between oropharyngeal and BAL taxa (e.g., *Atopobium*, *Haemophilus*, *Prevotella_7*, *TM7x*) were more frequent in FS6w than NS, partly overlapping with AS (Additional Figure 4).

**Figure 4.**
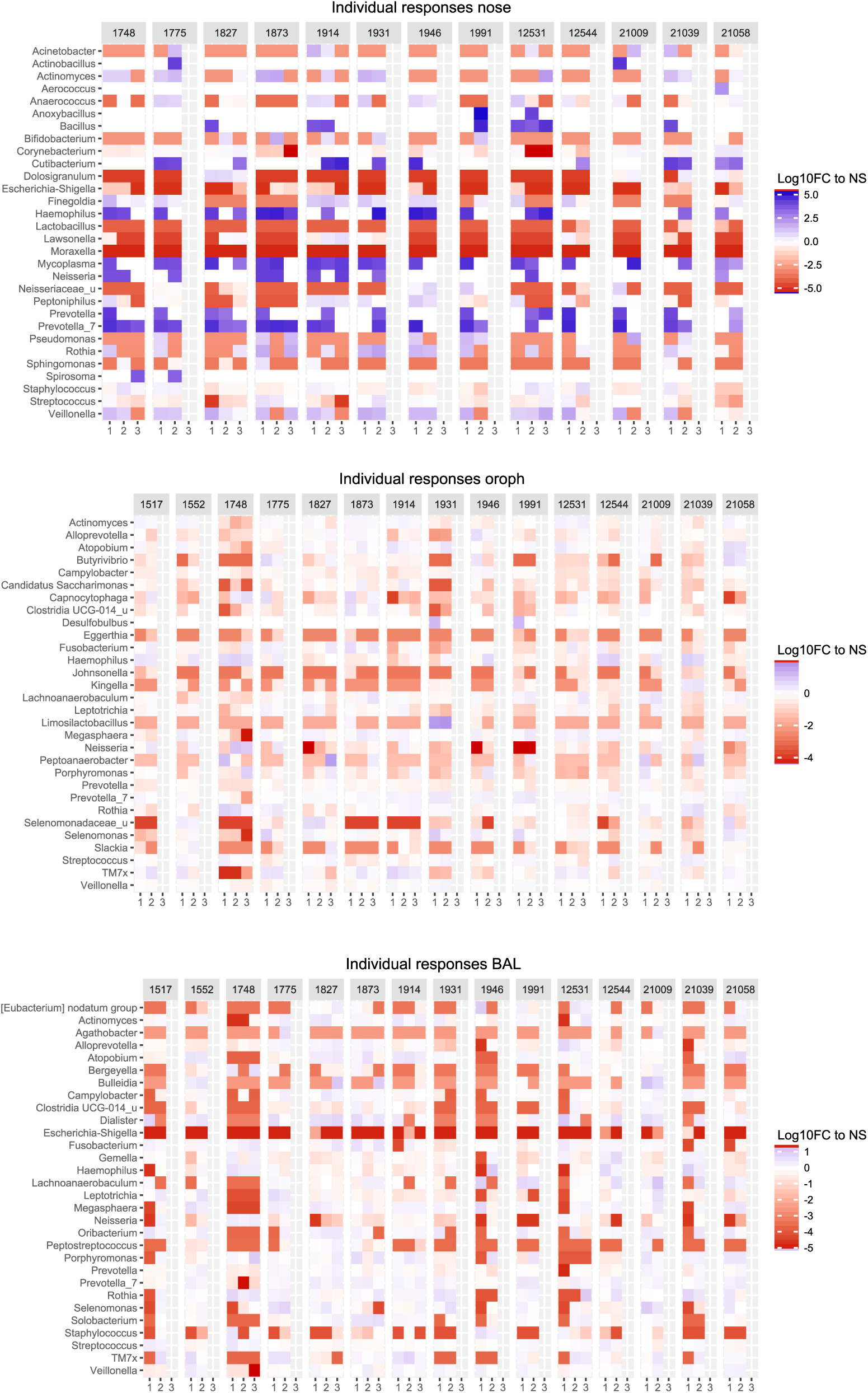
Individual responses to smoking and cessation Heatmaps show individual responses of taxa (the 10 most abundant genera as well as those genera showing the highest correlation to personalization and recovery, each 10) for different smokestatus (1: AS, 2: FS6w, 3: FS1yr) calculated as log-fold changes to NS baseline.

Personalization remained highest in the nose (0.87) and lower in the oropharynx (0.60) and BAL (0.62) (Fig. 3g). Compared to AS, variability increased in nose, remained stable in oropharynx, and decreased in BAL to NS levels. While recovery was generally impaired in heavy smokers, personalization of oropharynx and BAL remained positively associated with smoking history (Additional Figure 3).

### Long-term smoking cessation only leads to a partial and site-specific recovery

After 1 year of cessation (FS1yr), nasal richness remained elevated compared to NS, whereas BAL richness recovered to NS levels. Oropharyngeal richness increased relative to AS but was comparable to NS (Fig. 3a). However, microbial communities in all sites remained distinct from NS, with only a weak recovery trend in BAL (p=0.203) (Fig. 3b). Recovery potential of oropharynx and BAL remained negatively correlated with smoking years and pack-years, while nasal recovery showed strongest association with maximum cigarettes/day (Additional Figure 3).

Genus-level shifts reflected a partial recovery: in nose, *Haemophilus* and *Mycoplasma* decreased towards NS-level, whereas *Prevotella_7* remained elevated. In oropharynx, *Haemophilus*, *Mogibacterium*, *Parvimonas*, *Prevotella*, and *Prevotella_7* normalized, while *Porphyromonas* increased, and *Campylobacter* decreased compared to NS. In BAL, only *Bacteroides* recovered, while *Alistipes*, Muribaculaceae, and Prevotellaceae UCG-001 remained depleted, and *Parvimonas* and *Treponema* increased (Fig. 3c, Additional Figure 2).

Core microbiota analysis identified 2, 47, and 34 genera for nose, oropharynx, and BAL, respectively (Fig. 3e, Additional Table 2). FS1yr oropharynx harboured 12 unique genera, while BAL showed 11 FS1yr-specific genera and greater overlap with NS than AS (Fig. 3d, Additional Table 2). Across sites, only *Streptococcus* was shared universally, while 16 genera were oropharynx-specific and 2 were unique for BAL (Fig. 3e). Oropharynx–BAL overlap remained high, supported by continuously reduced between-group distances compared to NS (Fig. 3f). Taxa correlations between oropharynx and BAL persisted but were less pronounced than at 6 weeks (Additional Figure 4).

Personalization remained highest in nose (0.84) and lower in oropharynx (0.71) and BAL (0.69) (Fig. 3g). Compared to AS, variability increased in oropharynx, was stable in nose and BAL, but exceeded NS in all sites. Even after 1 year, personalization remained positively correlated with smoking history, highlighting the long-lasting imprint of smoking on respiratory microbiota. However, results should be interpreted with caution given the small sample size (n=5).

### Microbial response patterns are highly personalized

Beyond general smoking and cessation effects, high inter-individual variability was observed across all airway compartments (Fig. 4). In the nose, some subjects (e.g., 1991, 21058) exhibited persistently high personalization scores, with both shared and individual-specific responses to smoking and cessation. For instance, trends for several taxa were consistent across subjects (e.g. *Escherichia-Shigella*), whereas others varied markedly between individuals (e.g., *Neisseria*, *Dolosigranulum*, *Prevotella, Veillonella*).

In the oropharynx, personalization was generally lower, with subjects 1991 and 21058 showing only medium individuality, whereas others (1748, 1931) revealed high personalization. Smoking induced similar directional changes for several taxa (e.g. *Actinomyces, Alloprevotella, Prevotella*), but individual responses differed for specific genera (e.g. *Haemophilus, Neisseria*, *Prevotella_7*, *Rothia*, *Selemonas, Veillonella*), and these patterns persisted or shifted variably after cessation.

In BAL, highest personalization was seen in subjects 1748 and 1946. While general trends (e.g., reduced *Alloprevotella*, *Atopobium*, *Dialister*, *Prevotella*, *Selemonas*) were shared, responses of other taxa (e.g. *Actinomyces, Haemophilus*, *Neisseria*, *Rothia*, *Streptococcus*) were highly individual with personalization partially maintained after cessation.

Overall, personalization in the oropharynx and BAL positively correlated with the smoking history, whereas nasal microbiome individuality showed no clear association with smoking metrics (Additional Figure 3). Subjects with high nasal personalization displayed varying smoking histories (38 vs 8 smoking years and 40 vs. 8 pack-years for subjects 1991 and 21058, respectively), highlighting compartment-specific drivers. Concordant responses between oropharynx and BAL in highly personalized individuals, such as sustained increases in *Haemophilus* and *Rothia*, underscore the strong overlap between these sites and suggest microbial exchange along the respiratory tract.

### Microbial community assembly processes and network topologies

URT microbial community assembly was predominantly shaped by stochastic processes (Fig. 5). In nasal samples, dispersal limitation dominated in NS and FS1yr, while smoking slightly increased contributions of undominated drift and homogeneous selection, still persisting after 6 weeks cessation (FS6w). Oropharyngeal communities were largely shaped by dispersal limitation in NS, AS and FS6w, with undominated drift increasing after long-term cessation. BAL communities were influenced by both stochastic and deterministic processes, with homogeneous selection particularly prominent in AS and, to a lesser extent, in FS6w, while normalizing after 1-year cessation.

**Figure 5.**
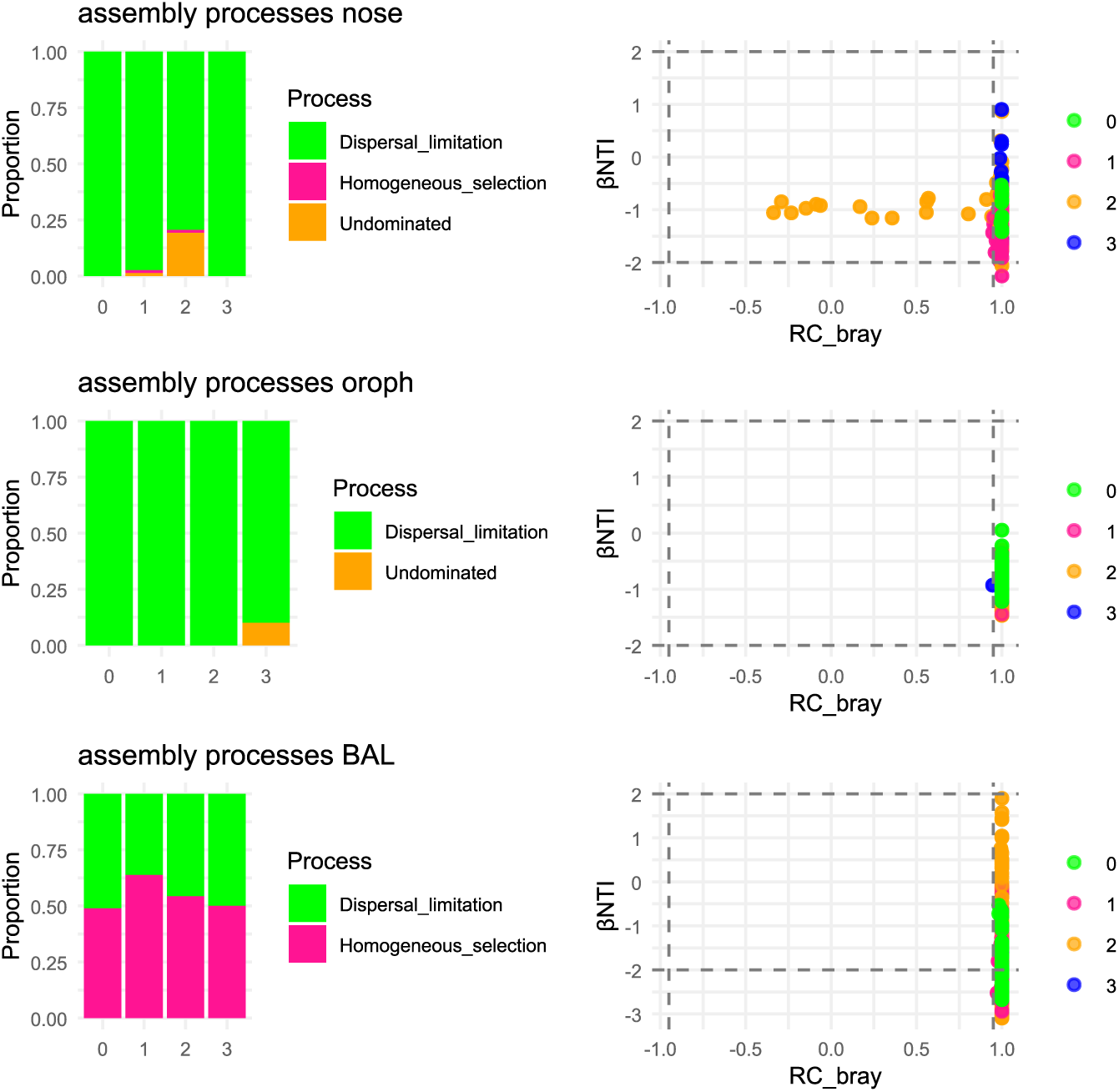
Community assembly processes dominating the different smokestatus in nose, oropharynx and BAL Differences in ßNTI and RCbray served to determine the processes shaping microbial communities. Analysis showed a dominance of stochastic processes at all stages ((0: NS, 1: AS, 2: FS6w, 3: FS1yr) in nose and oropharynx, whereas the BAL community was shaped by both stochastic and deterministic assembly, with temporarily higher contribution of selection in AS and after short-term cessation. The threshods for |βNTI| = 2 and |RCbray| = 0.95 are highlighted as horizontal and vertical dashed lines, respectively.

Microbial network analysis for the oropharynx and BAL revealed pronounced shifts during smoking and cessation (Fig. 6, Table 2). In oropharynx, smoking reduced network complexity (fewer nodes, lower edge density and degree centrality, reduced robustness) while increasing modularity and path length. Short-term cessation had minimal impact, whereas long-term cessation restored connectivity, robustness, and degree centrality, resulting in denser but less modular networks than in NS. In BAL, smoking induced stronger network reductions, with restructuring evident already at 6 weeks and continuing through 1 year, leading to networks with comparable robustness, edge density, and degree centrality, though modularity remained elevated relative to NS. Due to the low number of core ASV, nasal networks could not be calculated.

**Figure 6.**
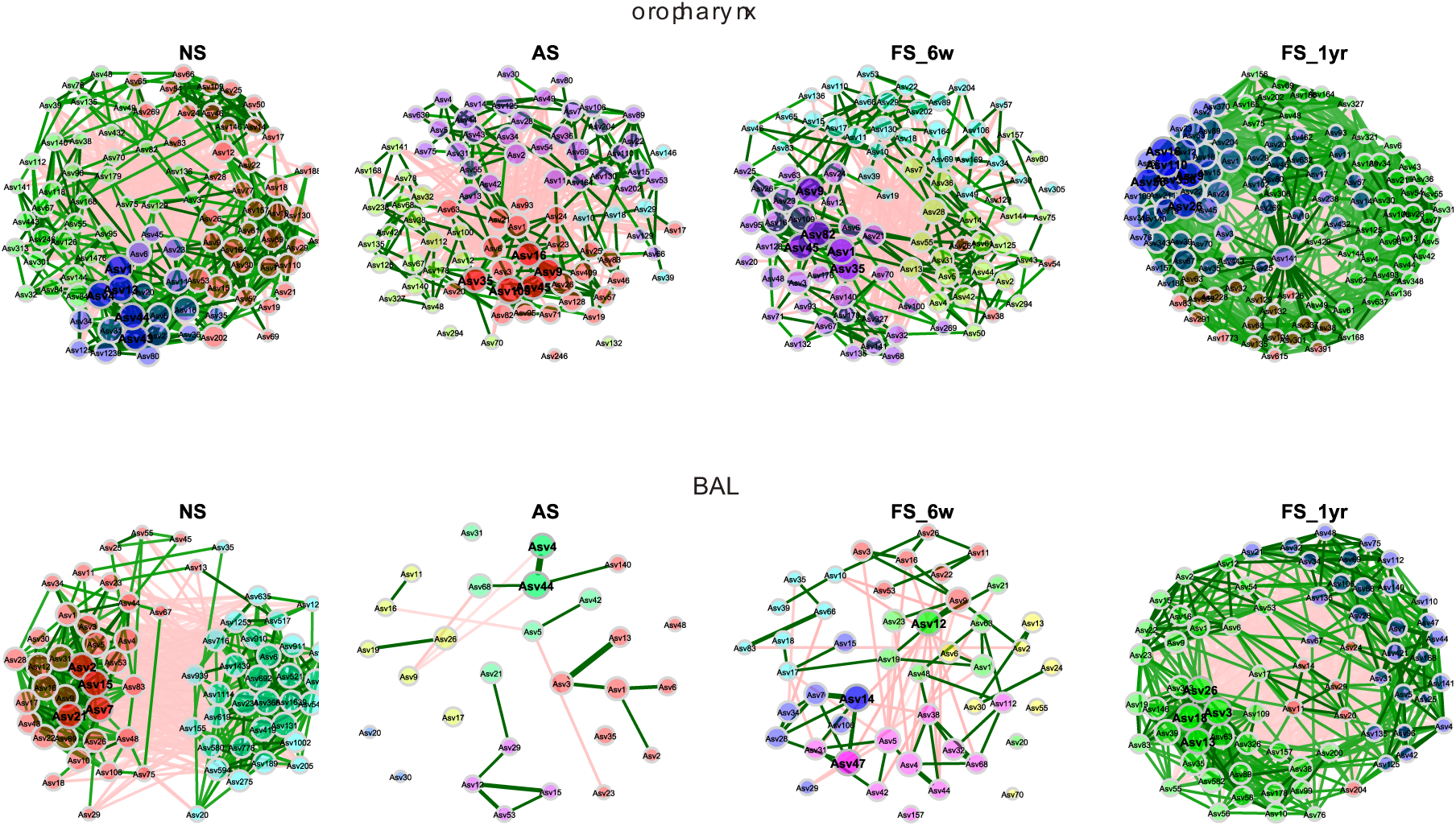
Microbial co-occurrence patterns among the different smokestatus Bacterial co-occurrence network analysis of core ASV (defined as ASV pesent in at least 50% of subjects) for NS, AS, and after smoking cessation (FS6w, FS1yr) in oropharynx and BAL. Nodes indicate ASV and edges represent Pearson correlation (threshold 0.3) between a pair of nodes. The green and red links represent positive and negative network interactions, respectively. Modules represent group of interconnected nodes, each differently colored. Bold nodes represent hub taxa, whereas node sizes indicate their respective eigenvectors. The table shows important topological features of bacterial co-occurrence networks, including number of nodes, clustering coefficient, modularity, positive edge percentage (PEP), edge density, robustness, average path length, centrality measures (degree, between, close, eigenvector) and number of modules.

## Discussion

Understanding how smoking perturbs the respiratory microbiome and whether these changes are reversible is critical for designing interventions to restore airway health. This longitudinal, multi-compartment design of our study nicely demonstrates that smoking induces site-specific alterations in the respiratory microbiome, with increased individuality in the nasal, oropharyngeal, and lung communities. Short-term cessation partially restores richness and community composition, particularly in the lower airways, whereas long-term cessation leads to only partial recovery and persistent personalization. Recovery trajectories are highly individualized, reflecting both stochastic and deterministic ecological processes and highlights the lasting impact of smoking on respiratory microbial network

### Microbial communities among the respiratory tract are niche-specific

Oropharyngeal and BAL microbiomes overlapped in NS, dominated by *Actinomyces, Prevotella* groups, *Streptococcus* and *Veillonella,* while nose was dominated by *Corynebacterium* and *Staphylococcus*. This reflects microaspiration and previous reports showing that the lung microbiome resembles the oropharynx, whereas the nose is closer to skin (^12, 40^). Each site hosts a unique, dynamic community shaped by airflow, mucociliary clearance, immune defences, and local microenvironments (^8, 41^).

### Effect of smoking and cessation on the microbiome is highly dependent on the airway cavity

Smoking alters airway microbiomes via oxygen depletion, antimicrobial activity, biofilm promotion, and immune impairment, enriching potential pathogens such as *Streptococcus pneumoniae* and *Haemophilus influenza* (^22, 42^). In our study, *Haemophilus* remained elevated in nose and oropharynx after 1 year cessation, linking smoking to COPD risk (^43, 44^). *Streptococcus* abundance was stable at the genus level, but ASV-level analysis revealed variable responses, emphasizing the importance of fine-scale resolution. Smoking enriched anaerobes such as *Prevotella* and *Veillonella,* which can modulate immunity and inhibit pathogens but may also increase susceptibility to inflammatory diseases if dysregulated (^45, 46^).

Oropharyngeal *Prevotella* responses were heterogeneous, reflecting intra-genus variability and oxygen sensitivity across clades. *Neisseria* decreased with smoking as reported previously (^12, 25^), likely due to oxygen depletion, but recovered rapidly post-cessation. Previously it was reported that *Neisseria* species could grow in anaerobic biofilms exhibiting denitrification (^47, 48^) and, by this, produce nitrite and possibly nitric oxide, which both may act as antimicrobial compounds. This may explain the fast re-establishment of *Neisseria* observed after cessation. Despite high shared genera and substantial resilience, oropharyngeal communities did not fully revert to that of NS after 1 year. Long-term cessation favoured colonization by biofilm-forming oral bacteria often associated with periodontitis like *Aggregatibacter, Dialister* and *Fretibacterium* (^49–51^), occupying niches freed by smoking-induced disturbance.

Smoking effects were even more pronounced in the lung. While bacterial diversity recovered after 1 year, community composition remained altered. *Alistipes, Bacteroides, Muribaculaceae*, and *Prevotellaceae* were depleted, likely disrupting immune and metabolic homeostasis. Oral- and gut-associated biofilm-forming taxa (e.g. *Capnocytophaga, Granulicatella, Parvimonas*) colonized the lung post-smoking cessation, reflecting URT-LRT microbial exchange and re-shaping of communities towards greater oropharynx-lung similarity. Our longitudinal data confirm that smoking imposes strong, lasting effects on airway microbiomes, with partial recovery over time.

### Smoking favours personalization long-lasting as result of site-specific assembly processes

Smoking increased individual variability across all airway sites, which persisted after cessation. In the nose, microbial community assembly was largely driven by stochastic processes, likely amplified by high environmental exposure (fluctuating temperature, humidity and airborne particles) and smoke-induced host effects, explaining high personalization. In contrast, the oropharyngeal microbiome, buffered by saliva (^52^), maintained a relatively stable core community but showed increased personalization associated with smoking duration and intensity, reflecting priority effects and niche colonization. Lung communities were strongly shaped by deterministic processes, consistent with its low microbial load, restricted immigration, and higher immune surveillance (^40, 53, 54^); smoking intensified host-driven selection, favouring specific taxa. Partial recovery post-cessation indicates that deterministic and stochastic forces jointly drive long-term individual patterns.

### Effect of smoking and cessation on microbial co-occurrence is dependent on the airway cavity

Smoking reduced the network connectivity, robustness, and centrality while increasing modularity and path length in oropharynx and BAL, reflecting disrupted microbial interactions. In the oropharynx, networks recovered slowly, eventually forming denser, less modular communities post-smoking cessation, indicating a reorganization of hub taxa and cooperative behaviour. In BAL, smoking-induced network reduction was more pronounced, but short-term cessation rapidly improved connectivity and robustness, continuing with long-term cessation. Persistent differences in hub taxa and modularity suggest lasting impacts of smoking, though functional recovery is substantial. Differences between oropharynx and BAL reflect ecological contrasts: high-dispersal, stochastic environments in oropharynx versus low-biomass, deterministic filtering in lung, shaping both intensity and speed of recovery.

### Strength and limitations of the study

Despite the strength of the study by analyzing the microbial communities among the entire respiratory tract and the longitudinal study design that enabled us to follow changes over time within the same individuals both before and after smoking cessation, this study has two main limitations: On the one hand, it is based on a relatively small number of participants and was mono-centric, thus results must not be necessarily generalizable to a larger, more diverse population. On the other hand, the used 16S rRNA gene-based metabarcoding approach is not suitable to explore the functional attributes of the identified microbial communities, which would be necessary to better understand microbial interactions. Thus, further longitudinal studies including long-read metagenome sequencing in combination with metabolomics are warranted to identify changes in the functionality of microbial responders and the overall community. Furthermore, such studies should be conducted with multi-centric design with respect to population heterogeneity.

## Conclusion

In conclusion, our study reinforces growing evidence that lungs harbor a diverse and dynamic microbiome essential for human health. In healthy individuals, their composition is mostly shaped by a balance of immigration into and elimination from airways, whereas altered growth conditions predominate during disease. Our findings highlight that (I) microbial communities among the respiratory tract are site-specific, with high overlap between oropharynx and the lower respiratory tract, (II) the response to cigarette smoke and cessation is largely site-dependent, most pronounced in the nasal and lung microbiomes, and (III) smoking increases individual variability in a site-specific manner through different community assembly processes, with effects still visible after long-term cessation. Furthermore, the high microbial exchange potential among different respiratory regions, resulting in gradual immigration of bacteria from the oral cavity or URT into “deeper” regions after reduction of smoking-induced stressors, underlines the need to analyze the entire respiratory tract microbiome rather than isolated compartments. to better understand of smoking and cessation effects.

Our findings could help explain the heterogeneity of smoking-related airway diseases (^55–57^): stochastic colonization in the upper airways could promote variable susceptibility to infections such as rhinitis or sinusitis, whereas deterministic enrichment of smoke-tolerant taxa in the lung may drive more consistent risks for COPD or asthma exacerbations.

Although partial microbial recovery after cessation suggests potential for therapeutic intervention, persistent alterations and highly personalized responses pose major challenges for developing universal strategies: (I) even after 1 year the respiratory tract microbiome of former smokers was not completely restored, indicating some non-reversible effects, and (II) the highly personalized response pattern driven by different assembly processes complicates the design of develop a generalized concept for interventions. For nose and oropharynx, strategies enhancing competitive colonization or microbial reseeding may o counteract stochastic divergence, whereas in the lung therapies focusing on restoring host environment and selective pressures may be more effective. However, for the development of targeted interventions to accelerate microbial recovery post-cessation, a deeper understanding of the compartment-specific balance between stochastic and deterministic forces driving the personalized smoking-related changes and recovery is necessary. Thus, future longitudinal, multi-center studies employing long-read metagenomics and metabolomics are needed to elucidate compartment-specific dynamics and functional implications. Nevertheless, this study provides valuable insights into the complex interplay between smoking, the airway microbiome and respiratory health, indicating that microbial imbalances may persist beyond cessation and contribute to long-term disease risk.

## List of Abbreviations

AS: active smokers;
ASV: amplicon sequence variants;
BAL: bronchoalveolar lavage;
COPD: chronic obstructive pulmonary disease;
cpd: cigarettes per day;
cpd_max: maximum cigarettes per day;
CRS: chronic rhinosinusitis;
FS: former smokers;
FS1yr: former smokers after 1 year cessation;
FS6w: former smokers after 6 weeks cessation;
GLMM: generalized linear mixed-effect model;
LRT: lower respiratory tract;
NMDS: non-metric multidimensional scaling;
NS: never-smokers;
RT: respiratory tract;
URT: upper respiratory tract;

## Consent for publication

Not applicable

## Availability of data and material

The sequence data generated in this study have been deposited in the NCBI Short Read Archive (SRA) under accession number PRJNA1328433 and will be made publicly available upon acceptance of the manuscript.

## Competing interests

The authors declare that they have no competing interests.

## Funding

Leibniz Competition 2016, ‘The lung microbiota at the interface between airway epithelium and environment’. The BioMaterialBank Nord is supported by the German Centre for Lung Research and member of popgen 2.0 network, which is supported by a grant from the German Ministry for Education and Research (grant number: 01EY1103).

## Acknowledgements

We would like to thank Susanne Walch, Cornelia Galonska, and Michaela Blank from the Research Unit Comparative Microbiome Analysis for their assistance with lab work and sequencing. Furthermore, the authors thank Andrea Glaewe, Johanna Döhling, Lenka Krabbe, Vanessa Schümann, Gabi Huss, Birgit Kullmann and Steffi Fox (all Research Centre Borstel) for coordinating and sampling the study participants and collecting clinical data.

This manuscript is a preprint and has not been peer reviewed.

## Authors contributions

S.G. took responsibility for data integrity, data analysis and drafting the manuscript. M.S., S.K-E., D.K., C.H., K.I.G., and J.O. contributed substantially to the study design and the revision of the manuscript. C.H. recruited the study subjects and obtained the microbiological samples.

## Preprint notice

This manuscript is a preprint and has not been peer reviewed.

## Additional files

**Additional Table 1.**
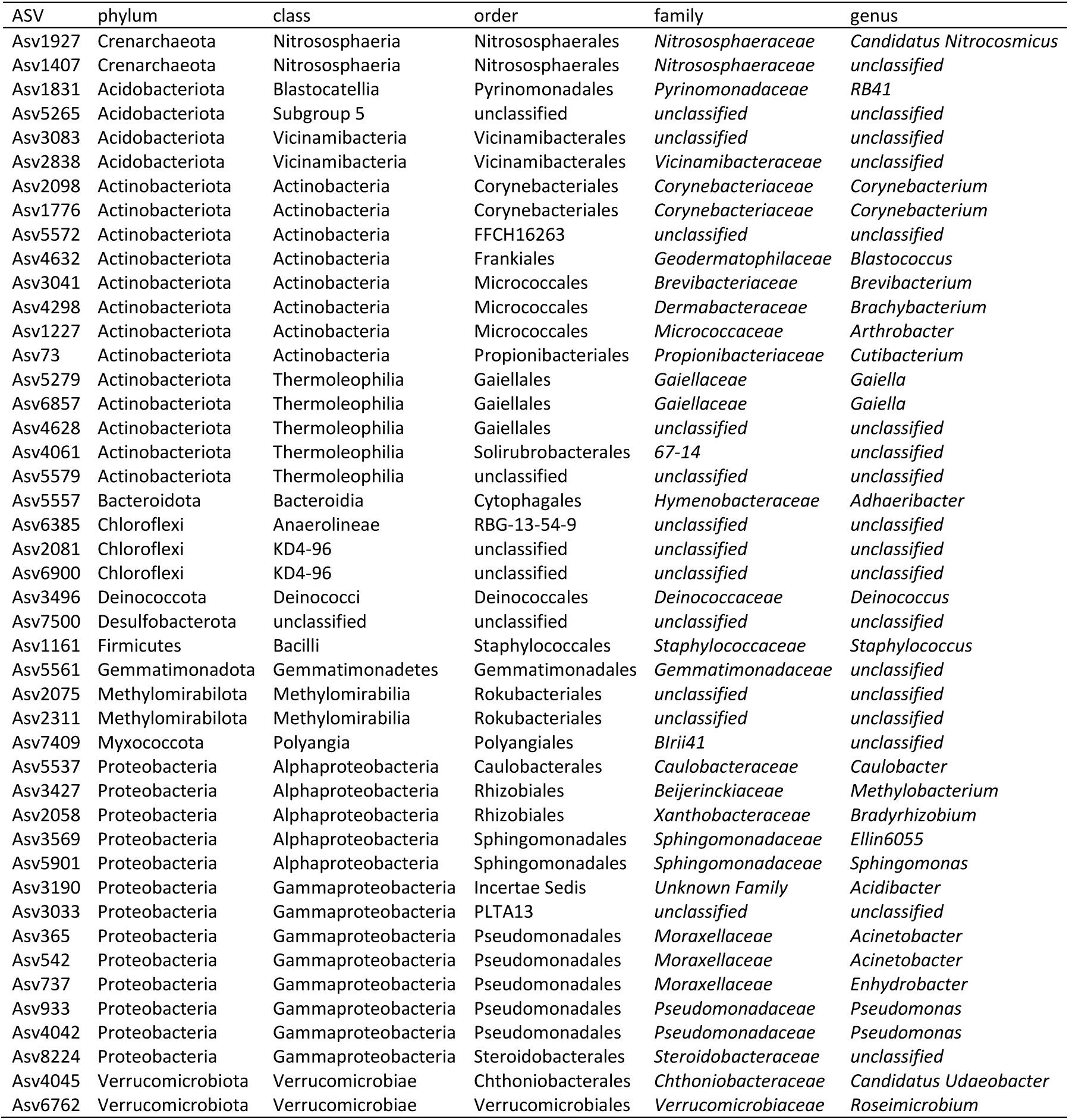
ASV identified as contaminants by core analysis and/or decontam were removed from data.

**Additional Table 2.**
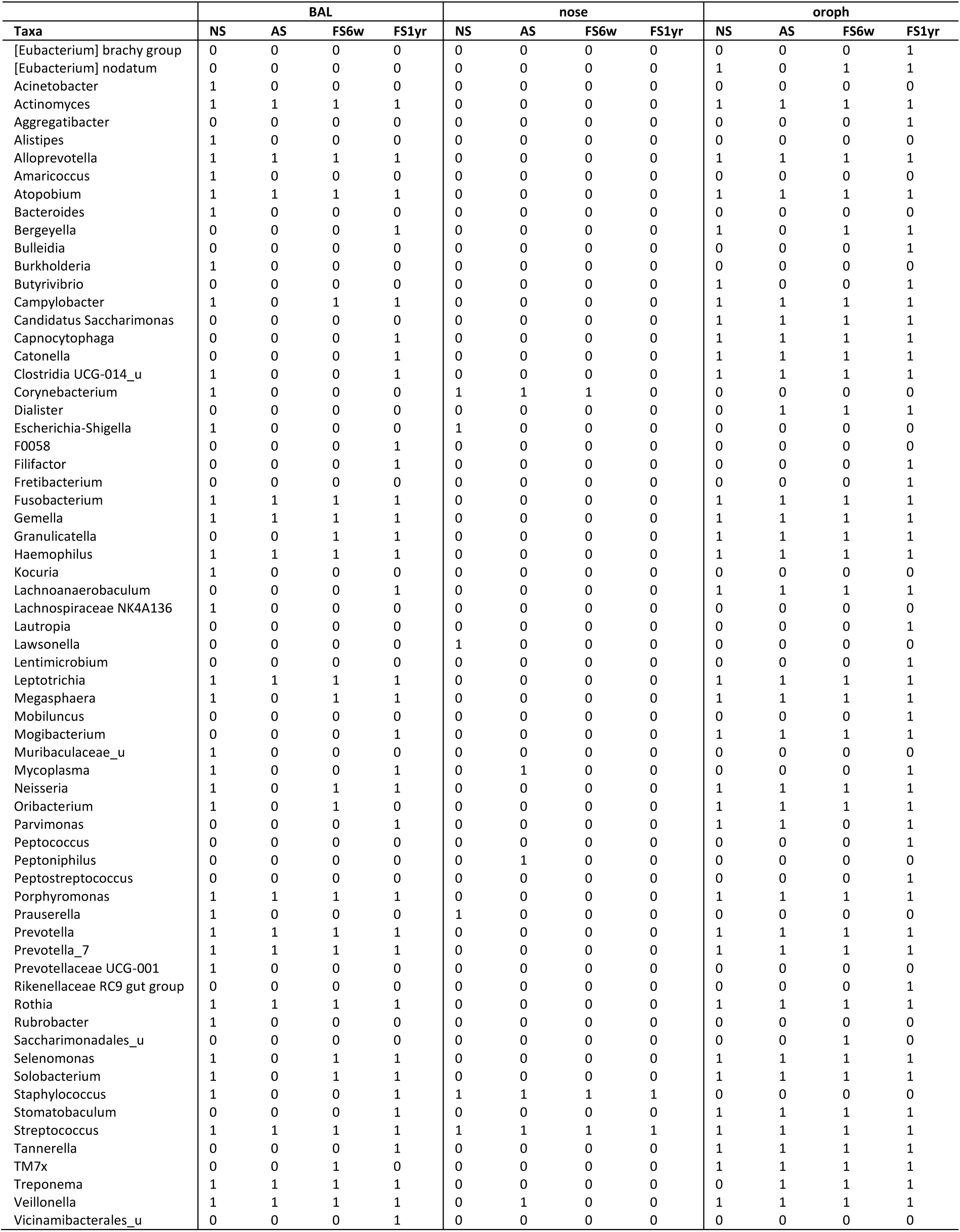
Core microbiota defined as genera observed in at least 80% of subjects per material (nose, oropharynx and BAL) and smoking status, shown as presence (1) and absence (0).

**Additional Figure 1.**
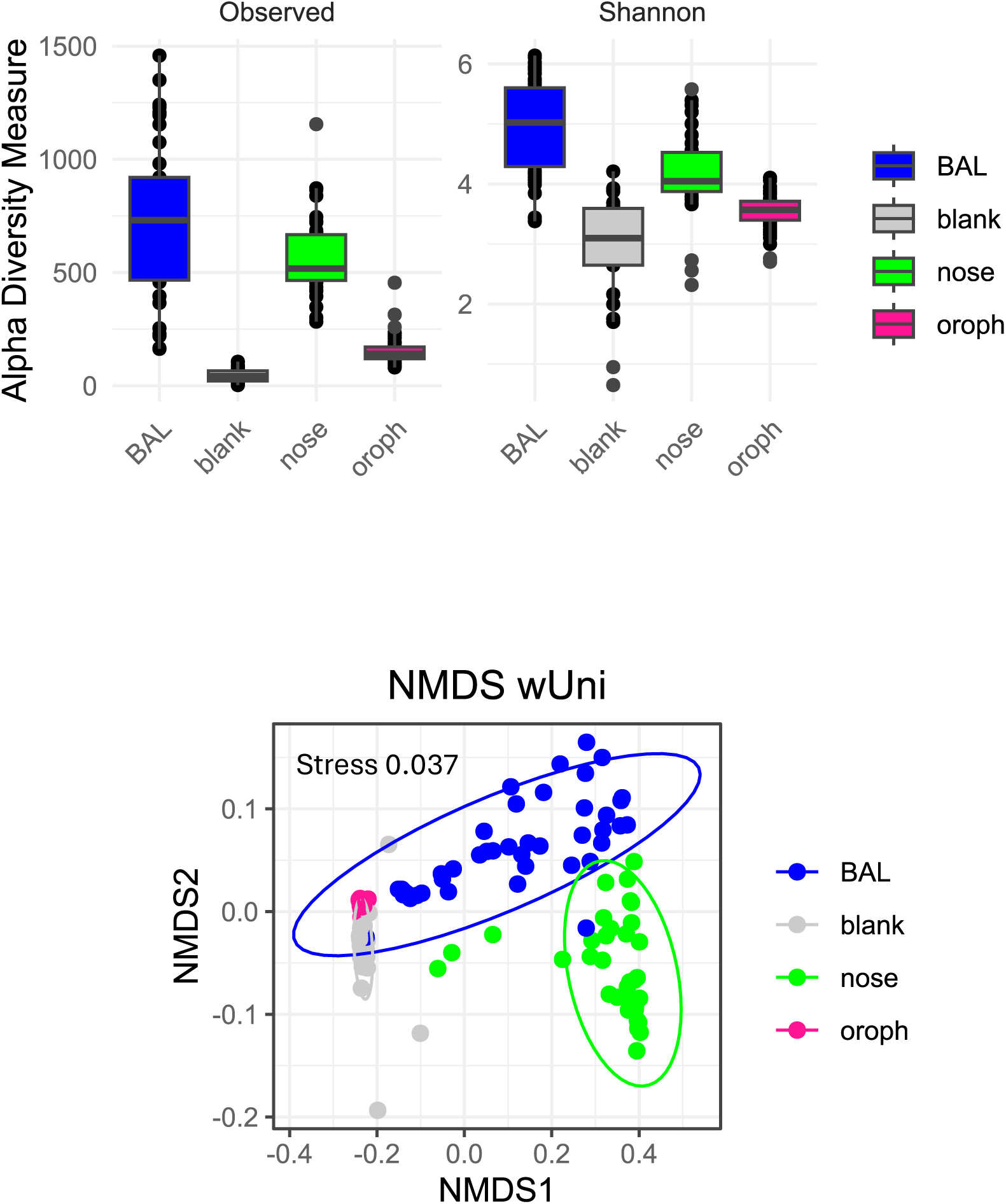
Alpha and beta diversity of respiratory tract samples showed a significantly higher diversity and different community composition compared to blank samples.

**Additional Figure 2.**
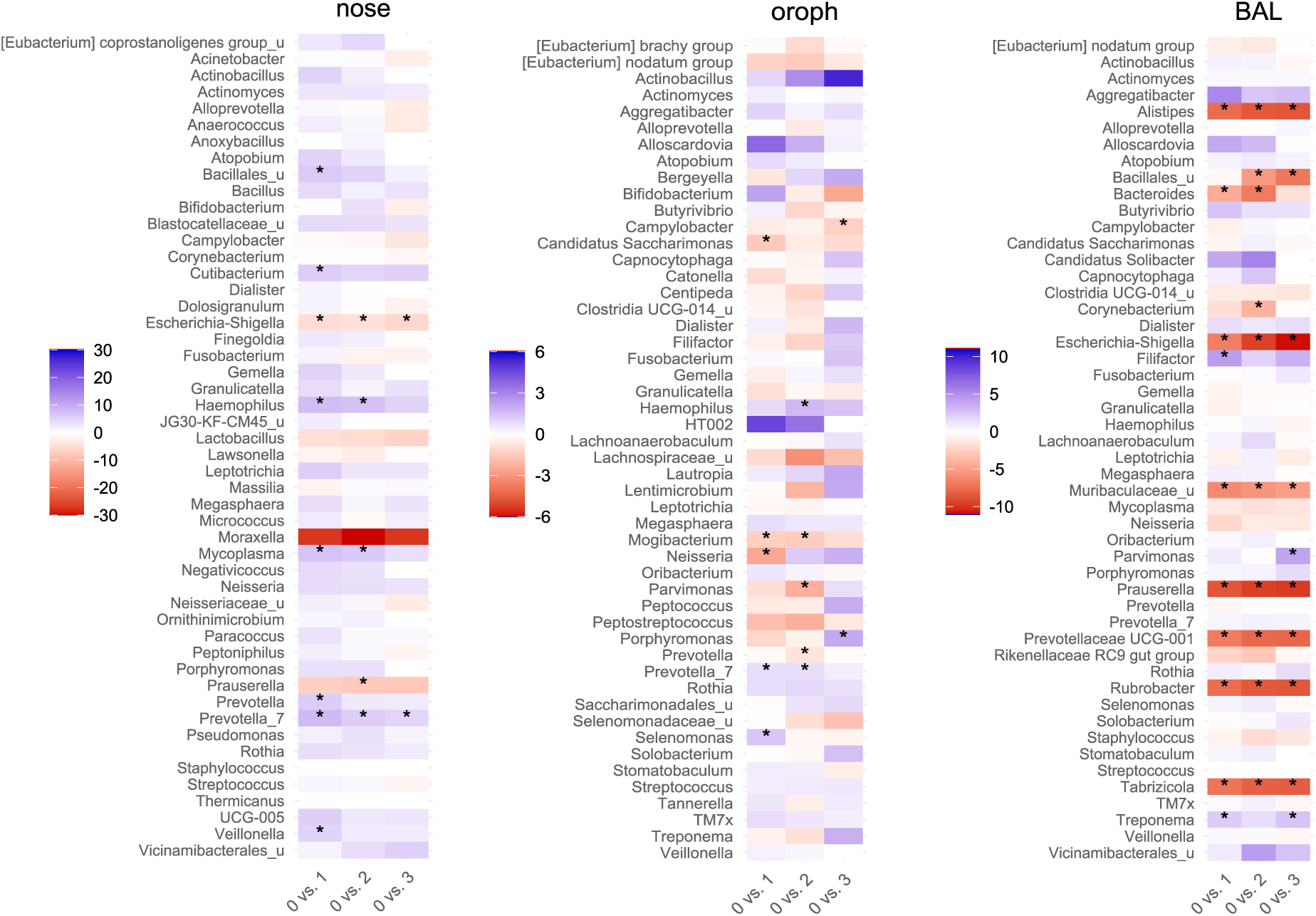
Heatmaps show changes of the top 30 taxa in nose, oropharynx and BAL for the different smokestatus (1: AS, 2: FS6w, 3: FS1yr), calculated as log-fold changes to NS baseline (0). Asterisks mark significant differences (p<0.05).

**Additional Figure 3.**
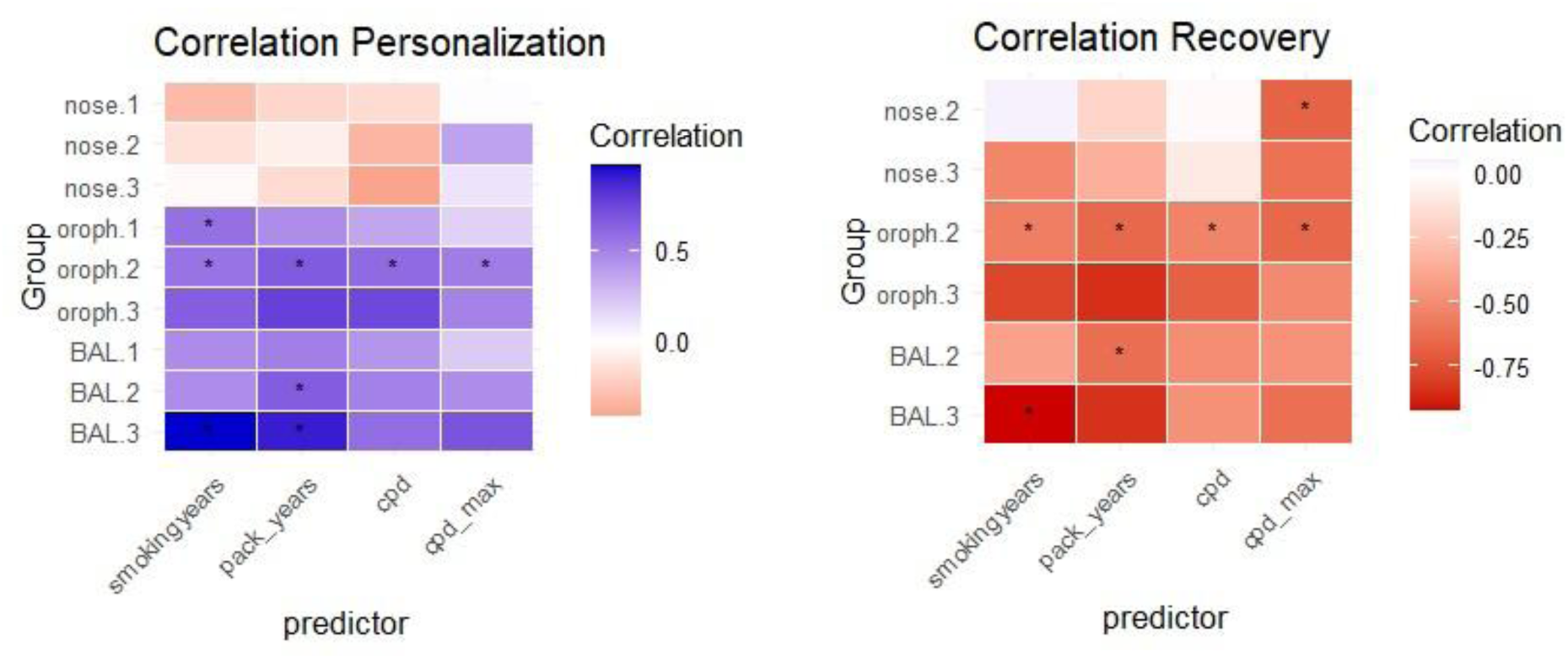
Correlations of personalization scores and recovery of microbial community with smoking-related factors (smoking years, packyears, cigarettes per day (cpd) and maximum cigarettes per day (cpd_max) for nose, oropharynx and BAL at the at the different smokestatus (1: AS, 2: FS6w, 3: FS1yr). Asterisks mark significant differences (p<0.05).

**Additional Figure 4.**
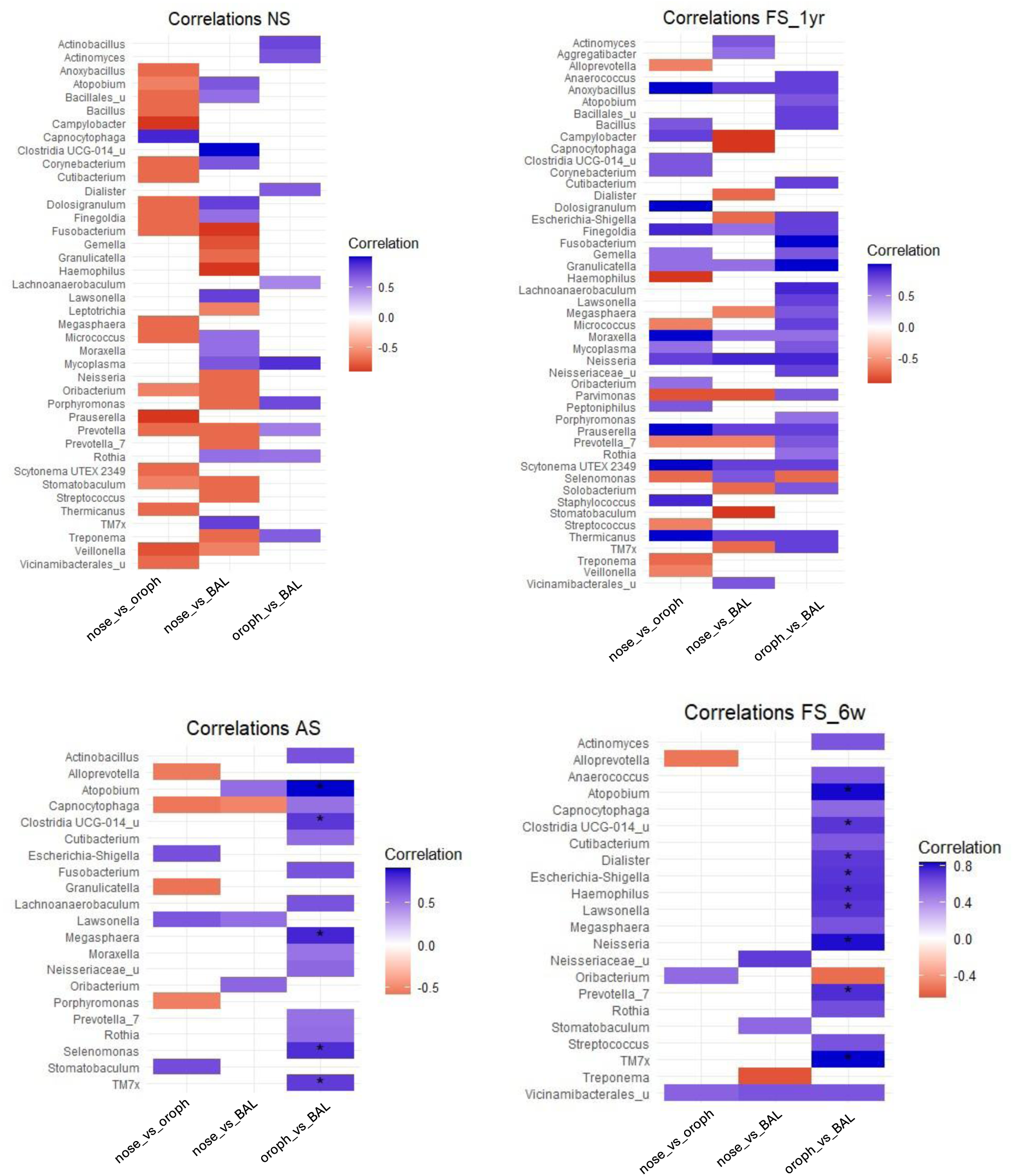
Correlations of taxa between the different respiratory sites in NS, AS, FS6w and FS1yr. Shown are all correlations > 0.5. Asterisks mark significant differences (p<0.05).

## Notes

### Competing Interest Statement

The authors have declared no competing interest.

